# Proteomic and phosphoproteomic analysis identifies novel liver-related signaling in retinal pigment epithelial cells during epithelial-mesenchymal transition

**DOI:** 10.1101/2020.02.17.953315

**Authors:** Joseph L. Mertz, Srinivas Sripathi, Xue Yang, Lijun Chen, Noriko Esumi, Hui Zhang, Donald J. Zack

## Abstract

Epithelial–mesenchymal transition (EMT) of the retinal pigment epithelium (RPE) is associated with several potentially blinding retinal diseases. Proteomic and phosphoproteomic studies were performed on human pluripotent stem cell-derived RPE (hPSC-RPE) monolayers to better understand the pathways mediating RPE EMT. EMT was induced by enzymatic dissociation or by co-treatment with transforming growth factor beta (TGFβ) and tumor necrosis factor alpha (TNFα; TGNF). The global and phosphoproteomes were analyzed at 1 hr post EMT induction to capture early events in kinase/phosphatase signaling cascades and at 12 hrs to define early changes in protein abundance. Pathway enrichment analysis revealed that TGNF and Dissociation rapidly perturbed signaling in many of the same pathways, with striking similarity in the phosphoproteome at 1 hr. Surprisingly, functions related to liver cell proliferation and hyperplasia were strongly enriched in the phosphosites altered by both treatments at 1 hr and in protein abundance changes at 12 hrs. Hepatocyte Growth Factor-cMET signaling exhibited the strongest overall enrichment in both treatments. These signaling pathways may serve as suitable targets for the development of therapeutic strategies for the inhibition of RPE EMT, and thus progression of several debilitating visual diseases.

## INTRODUCTION

Disorders of the retinal pigment epithelium (RPE) cause blindness and diminish vision for millions of people around the world. In age-related macular degeneration (AMD), the leading cause of vision loss in developed countries, atrophy, mottling, neovascularization, and drusen deposits in the RPE are precursors to vision loss (1), while proliferative vitreoretinopathy (PVR) is characterized by RPE cell migration and proliferation (2,3). The healthy mature RPE is a monolayer of epithelial cells between the neural retina and vascular choroid (4), with microvilli that envelop photoreceptor (PR) outer segments (5), basement membranes that abut the choroid, and tight junctions between each other that provide an important part of the blood-retinal barrier (6). The RPE largely functions to support PRs: it conveys nutrients toward (7) and waste products away from the highly metabolic PRs (8), maintains ion concentrations for phototransduction (9), and phagocytoses the rapidly turned over PR outer segments to eliminate toxic byproducts (10). Particularly important, the RPE partners with the PRs to carry out the visual cycle – RPE cells take up the opsin chromophore all-trans retinal from PRs, isomerize it back to 11-cis retinal, and then export it back to the PRs (11). It also absorbs excess light via its abundant melanin pigmentation (12,13).

Epithelial-mesenchymal transition (EMT) is an important biological process in which epithelial cells lose their intercellular junctions, cellular polarity, and become more migratory. While EMT is integral to healthy embryogenesis and tissue morphogenesis, EMT can also play a role in pathology, for example by contributing to unwanted fibrosis and tumor metastasis. During RPE EMT, cells lose their cobblestone morphology and apical-basal polarity, and adopt the spindle-shape morphology and increased proliferation and migration properties of mesenchymal cells (14). This process contributes to several retinal diseases. In AMD, for example, long lasting structural disruption and sustained TGFβ signaling can cause RPE cells to transition to a more mesenchymal phenotype, which can be associated with subretinal fibrosis and can enhance disease progression (15,16). In patients with PVR, exposure to growth factors such as TGFβ and platelet-derived growth factor (PDGF) induces EMT in RPE cells, which in turn causes them to proliferate into epiretinal membranes that create longer term complications (17).

Many of the signaling pathways that initiate and maintain EMT act through protein phosphorylation and protein abundance changes. As kinase/phosphatase signaling can occur almost instantaneously in response to stimuli, changes in protein phosphorylation provide one of the earliest signatures of EMT induction. Thus, phosphoproteomics provides a powerful approach to study and define cell signaling pathways related to the induction of RPE EMT. Additionally, detailed proteomic analysis can provide insights into how RPE cells respond following the induction of EMT. Here, using human pluripotent stem cell-derived RPE cells (hPSC-RPE), we describe the application of these complementary approaches to characterize the system-wide phosphorylation and protein expression changes that drive RPE EMT. Our results complement previous findings that implicate HGF-Met in the cell wound response and migratory behavior of the RPE, phenomena that are linked to EMT (18–20). EMT has been well characterized in systems-based examinations of other tissues and cell types (21–25), and target-based analyses of EMT have been conducted in RPE (26–29), but to date EMT has not been thoroughly characterized in RPE. It is our hope that our findings provide a useful resource for future study of a variety of RPE diseases.

## METHODS

### 1. Derivation and culture of RPE from hPSCs

Human iPSCs derived from IMR-90 fibroblasts (30) were cultured and differentiated into RPE as previously described (31). Briefly, the iPSCs were maintained in mTeSR1 media (Stem Cell Technologies), on growth factor reduced Matrigel substrate (BD Biosciences), under 10% CO2 and 5% O2, and amplified by clonal propagation using the ROCK inhibitor blebbistatin (Sigma). For differentiation, iPSCs were plated at 25,000 cells/cm^2^ and maintained in mTeSR1, which was replaced with differentiation media for 40-45 days. Differentiating cells were enzymatically dissociated using 0.25% (wt/vol) collagenase IV (Gibco) and resuspended in AccuMAX (Sigma-Aldrich) to create a single cell suspension, then re-plated on to fresh Matrigel coated plates. They were then maintained in RPE medium (70% DMEM, 30% Ham’s F12 nutrient mix, 2% B27-serum-free supplement, 1% antibiotic-antimycotic solution (Invitrogen)) for 2-3 months to form mature RPE monolayers. hPSC-RPE cells were harvested and protein was extracted after five duplicate treatment conditions: 1) untreated monolayer; 2) 1 hr combined treatment with 20 ng/mL TGFβ plus 20 ng/mL TNFα (TGNF); 3) 12 hr 20 ng/mL TGNF treatment; 4) 1 hr after enzymatic dissociation with AccuMAX and replating; and 5) 12 hr after enzymatic dissociation and replating.

### 2. Phosphoproteome profiling

Protein was harvested in urea ammonium bicarbonate lysis buffer with complete EDTA free protease inhibitor (Roche) and phosphatase inhibitor (Sigma Aldrich), reduced by DTT, alkylated by IAA, and proteolytically cleaved by trypsin (Promega). The ten samples were labeled by TMT 10-plex isobaric tags (ThermoFisher) and complete labeling assessed by LC-MS/MS. Samples were pooled in a 1:1 ratio and then separated into 96 fractions on a 250 mm C18 column via high-pH reverse-phase LC (Agilent) with a gradient shifting over the course of 120 minutes from ammonium formate and 2% ACN Buffer A to ammonium formate and 90% ACN buffer B. These 96 fractions were concatenated down to 24 in a pattern that maximized fraction complexity. 10% of these fractions was used for global proteome analysis, while the remaining 90% was concatenated down further to 13 fractions, then subjected to Fe3+ immobilized metal affinity chromatography for phosphopeptide extraction. Ni-NTA agarose beads (Qiagen) were bound to Fe^3+^ after stripping them of previously bound metal ions using EDTA and extensive rinsing. Peptide fractions were incubated with Fe^3+^ beads, unbound peptides removed by removing supernatant and rinsing, then the bead suspension was transferred to C18 stage tips. Within the C18 tips, phosphopeptides were eluted from the immobilized metal affinity chromatography (IMAC) beads and then the C18 resin, resulting in a final elution of desalted phosphopeptides.

#### 2.1. LC-MS/MS data collection

Global proteome and phosphoproteome samples were separated on a Dionex Ultimate 3000 RSLC nano system (Thermo Scientific) with a PepMap RSLC C18 column (75 μm x 50 cm, 2 μm; Thermo Scientific) protected by an Acclaim PepMap C18 column (100 μm x 2 cm, 5 μm; Thermo Scientific) before injection into a Q-Exactive mass spectrometer (ThermoFisher). The mobile phase consisted of Buffer A (0.1% FA, 3% ACN) and Buffer B (0.1% FA, 90% ACN) and flow rate was 0.300 μL/min. Global samples were eluted from the column using a gradient of 10 min 2 to 5% Buffer B, 80 min 5 to 30% B, 22 min 30 to 38% B, 3 min 38 to 95% B, 10 min 95% B, 4 min 95 to 2% B, and 1 min 2% B. Phospho samples were eluted by a gradient of 10 min 2 to 7% B, 80 min 7 to 26% B, 22 min 26 to 34% B, 3 min 34 to 95% B, 10 min 95% B, 4 min 95 to 4% B, and 1 min 4% B. Spray voltage was set to 1.5 kV for both sample sets. Full MS spectra were collected from 400–2000 m/z at a resolution of 70,000 with an AGC target of 3×10^6^ and maximum injection time 60 ms. Data-dependent HCD MS2 spectra were collected from the top 12 most abundant ions with a 1.4 m/z isolation window and dynamic exclusion of 30s. MS2 spectra were collected at 35,000 resolution from 200 to 2000 m/z and 110 m/z fixed first mass, with a 2×10^5^ AGC target, 120 ms maximum injection time, 32% NCE, 3.3 × 10^4^ intensity threshold, peptide match set to ‘preferred’, and rejection of 1 and >8 charge states. Peptide sequences were matched against the Uniprot human proteome database (released 7/17/18), quantified by reporter ion intensity, and assigned phosphosite probabilities using MaxQuant Version 1.6.5 (32). Data analyses were performed with the DEP R package (33), Perseus and Ingenuity Pathway Analysis software packages (IPA; Qiagen), in addition to Database for Annotation, Visualization and Integrated Discovery (DAVID) (NCI, (34)) functional annotation tools online, the STRING database (www.string-db.org), Cytoscape, PhosphositePlus (Cell Signaling Technology, (35)), and NetworKin & NetPhorest under the KinomeXplorer platform (Linding and Jensen, (36)).

## RESULTS

### Global and phosphoproteomics results

To characterize the early system-level protein expression and phosphorylation responses of hPSC-RPE cells to EMT induction, we performed global proteomic and phosphoproteomic analyses on hPSC-RPE monolayers in which EMT was induced by either: 1) co-treatment with TGFβ+TNFα (TGNF) or 2) enzymatic dissociation with AccuMAX. For both treatments, cells were analyzed at 1 and 12 hrs post-EMT induction. Both timepoints were analyzed in duplicate and compared to a reference duplicate of untreated monolayers. Proteins were extracted and prepared for bottom-up LC-MS/MS global and phosphoproteomic analysis (Fig 1). TMT-10 isobaric labeling was used to facilitate quantitative assessment of EMT induced protein alterations, and in order to increase coverage of the phosphoproteome, phosphopeptides were enriched by iron immobilized metal affinity chromatography (Fe-IMAC) and the eluate was analyzed in separate LC-MS/MS runs from the global proteome.

**Figure 1.**
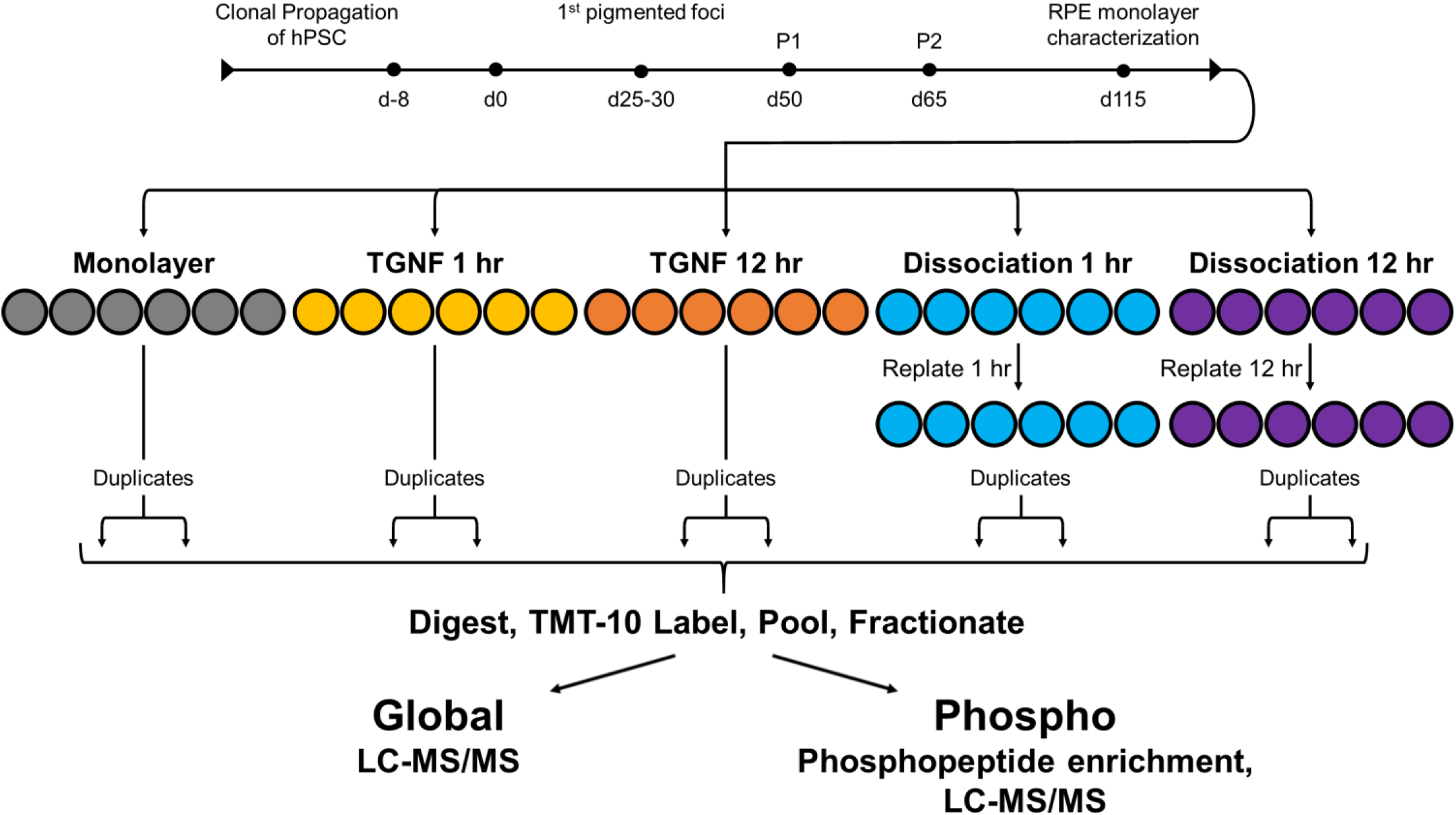
Global proteome and phosphoproteome workflow. After a 115 d differentiation procedure, duplicates of untreated hRPE monolayer served as control for comparison to duplicates of 4 treatment conditions: 1 hr combined treatment with 20 ng/mL TGFβ plus TNFα (TGNF); 12 hr 20 ng/mL TGNF treatment; 1 hr replating after enzymatic dissociation with AccuMAX; 12 hr replating after enzymatic dissociation.

8,648 total protein groups were quantified from the combined global and phosphoproteome analyses, with 7,606 matched to at least 2 unique peptides and 7,109 matched to at least 2 unique peptides in only the global fraction. We identified 15,346 phosphosites via the phosphoproteome workflow and 508 via the global workflow, with 383 of these overlapping. 9,338 phosphosites across 3,379 proteins were assigned ≥0.75 phosphate localization probability by MaxQuant software, thus meeting criteria for class I phosphosite designation. 943 or 10.2% of these class I sites were not reported in the Phosphosite Plus human dataset last modified on May 2, 2019 (35). The correlation coefficients for comparisons of reporter ion intensity between replicates (all greater than 0.98) as well as fold-change over control between replicates (ranging from 0.79 to 0.92) showed high reproducibility for both global (Fig 2) and phosphosite data (Fig 3). We also compared the protein abundance changes 12 hrs after dissociation to a previous analysis with similar parameters and found a high degree of correlation (Fig 4D).

**Figure 2.**
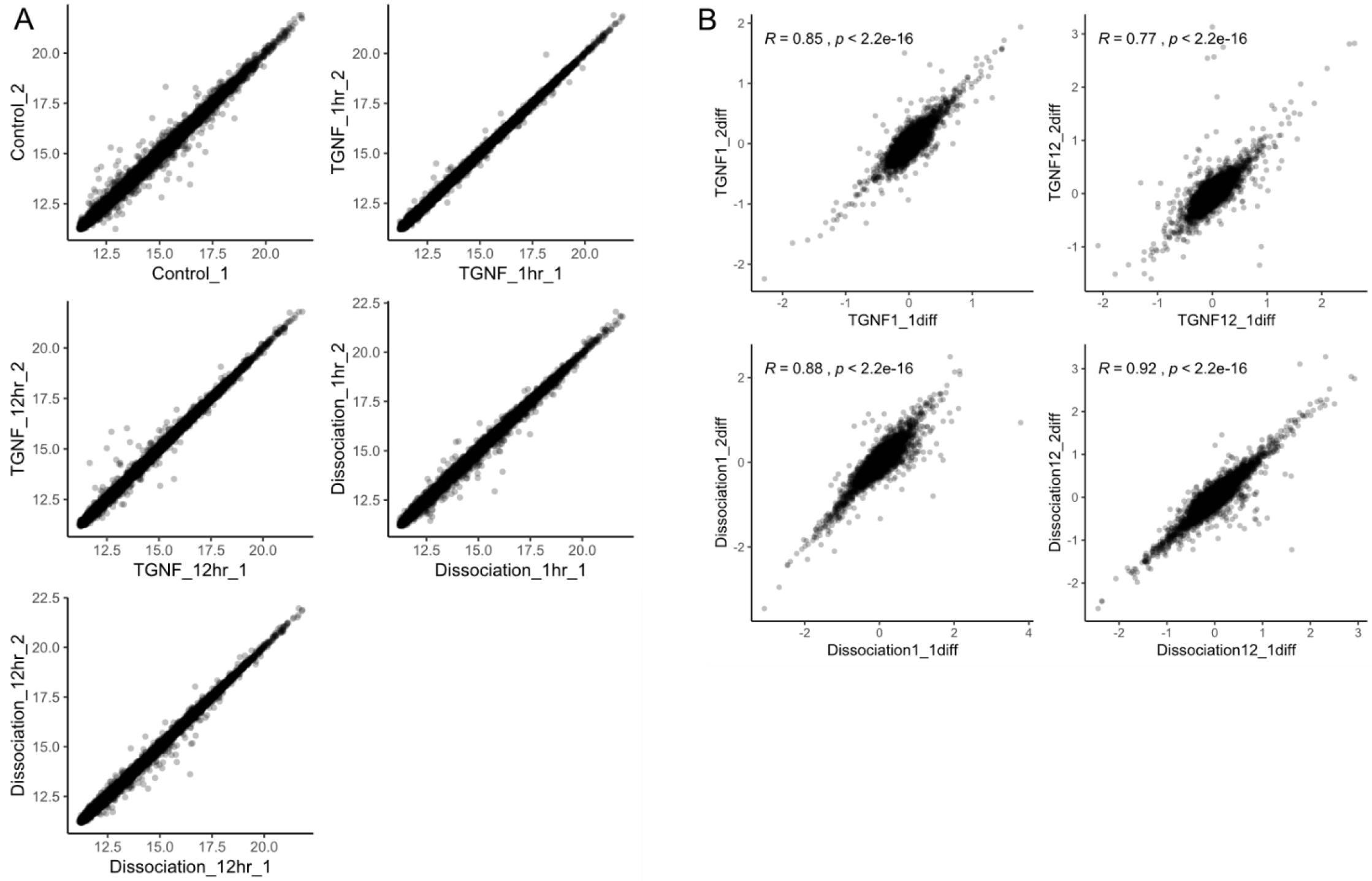
Global proteome data exhibit high reproducibility. A) Scatter plots comparing protein reporter ion intensities from duplicates of each treatment show very high correlation across the proteome. B) Comparison of treatment duplicate protein intensity changes from mean control values exhibits high correlation.

**Figure 3.**
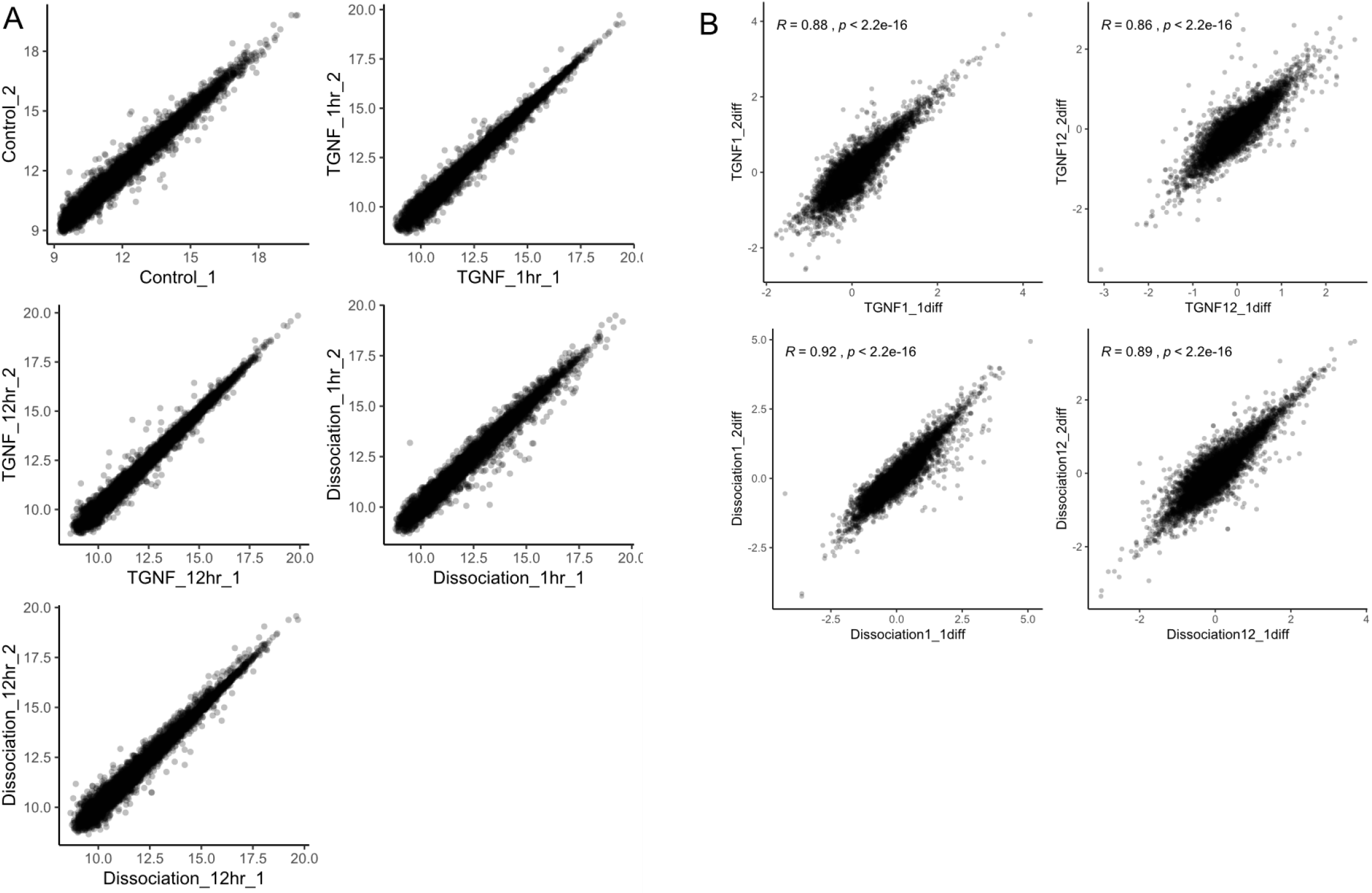
Phosphoproteome data exhibit high reproducibility. A) Scatter plots comparing phosphopeptide reporter ion intensities from duplicates of each treatment show very high correlation across the phosphoproteome. B) Comparison of treatment duplicate phosphopeptide intensity changes from mean control values exhibits high correlation.

**Figure 4.**
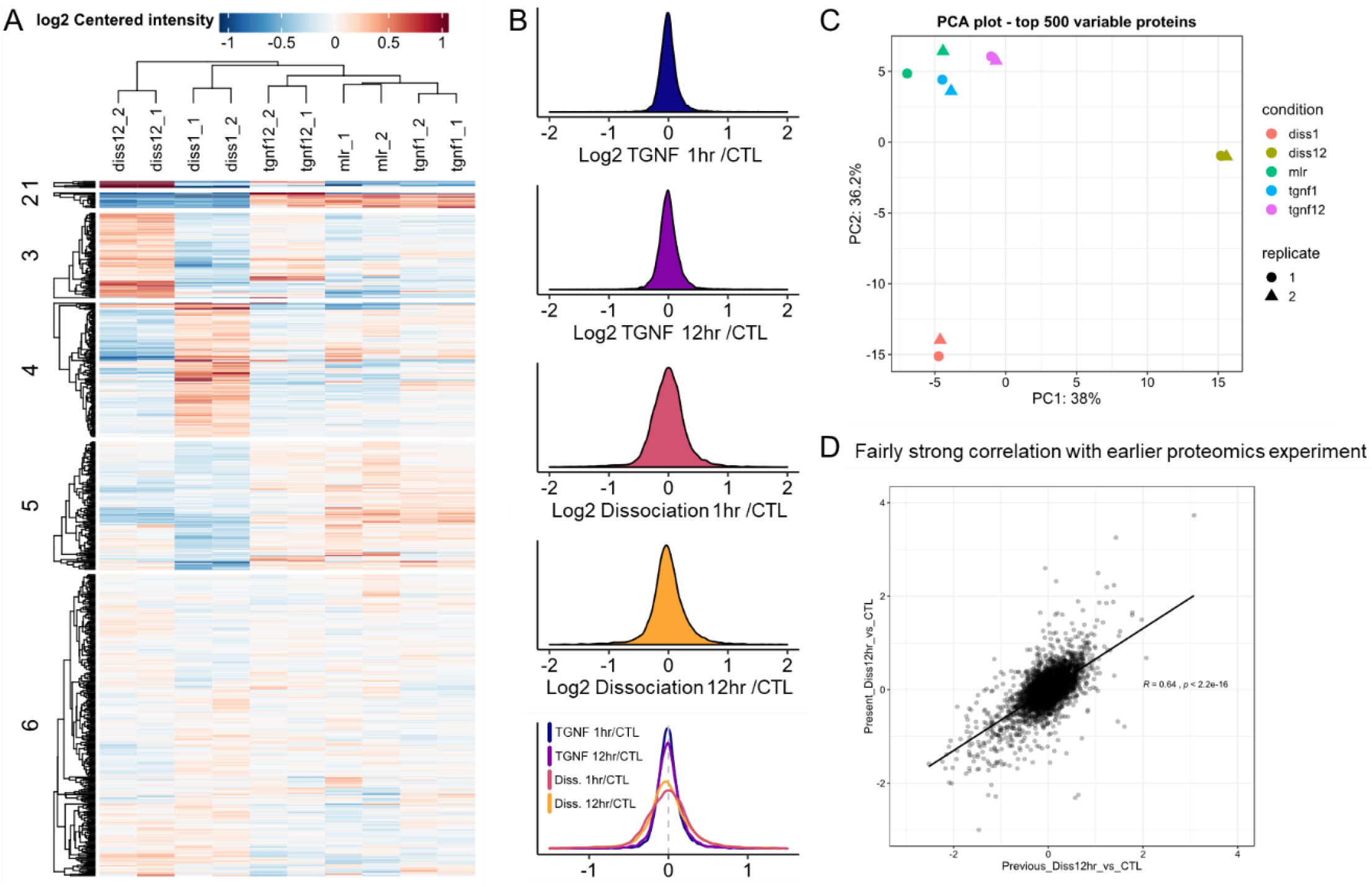
Enzymatic dissociation elicits more changes upon the proteome than TGNF treatment. A) Hierarchical clustering of global proteome data places replicates of each treatment most closely to each other and the two TGNF time points (1 and 12 hrs) most closely to untreated monolayer, followed by 1 and 12 hr dissociation, respectively. B) Histograms of protein changes over control for each treatment show more gross changes are caused by dissociation. C) Principle component analysis also shows tight clustering of duplicate samples, with less commonality between the dissociation samples and untreated control. D) Correlation analysis of proteome datasets from two separate 12 hr dissociation experiments performed more than a year apart shows high correlation in protein changes over control.

To determine significantly altered proteins and phosphosites we compared each treatment to control via student’s T-test and used a Benjamini-Hochberg adjusted *p*-value cut-off of 0.05 combined with a log_2_ fold-change (log2FC) threshold of 0.5 (Fig 5). Phosphosite fold-changes were normalized to their respective protein abundance changes before these comparisons to prevent changes in protein levels from enhancing or suppressing observed phosphorylation changes. Enzymatic dissociation induced more widespread protein abundance changes than TGNF treatment (561 vs 199), and both treatments induced more such changes at 12 hrs than at 1 hr (Diss: 628 vs. 258; TGNF: 116 vs. 106) (Fig 6A). Very few 1 hr TGNF protein changes persisted to 12 hrs (23 of 106), while almost half of protein changes caused by dissociated persisted (125 of 258). TGNF induced more phosphorylation changes than dissociation at 1 hr (513 vs. 395), but those induced by dissociation appeared to be more salient. The vast majority of 1 hr TGNF phospho changes were not altered at 12 hrs (364 vs. 149) with few phosphosites significantly altered only at 12 hrs (112). With dissociation, roughly equal proportions of phosphosites were observed only at 1 hr, only at 12 hrs, or both (222, 249 and 173, respectively). These trends can also be observed when examining the change over time of all altered proteins and phosphosites (Fig 6D). When comparing similarities in the changes induced by these two treatments, aside from phosphosite changes at 1 hr, most comparisons showed roughly half of TGNF induced changes overlap with dissociation induced changes, with rather small proportions of the corresponding dissociation induced changes overlapping (Fig 6B). This is not true for phosphorylation changes at 1 hr, however, as a large majority of the dissociation induced changes were also induced by TGNF. This comparison exhibited the strongest overlap of any, including those between different timepoints of the same treatment, suggesting the two treatments trigger a shared core of signaling responses.

**Figure 5.**
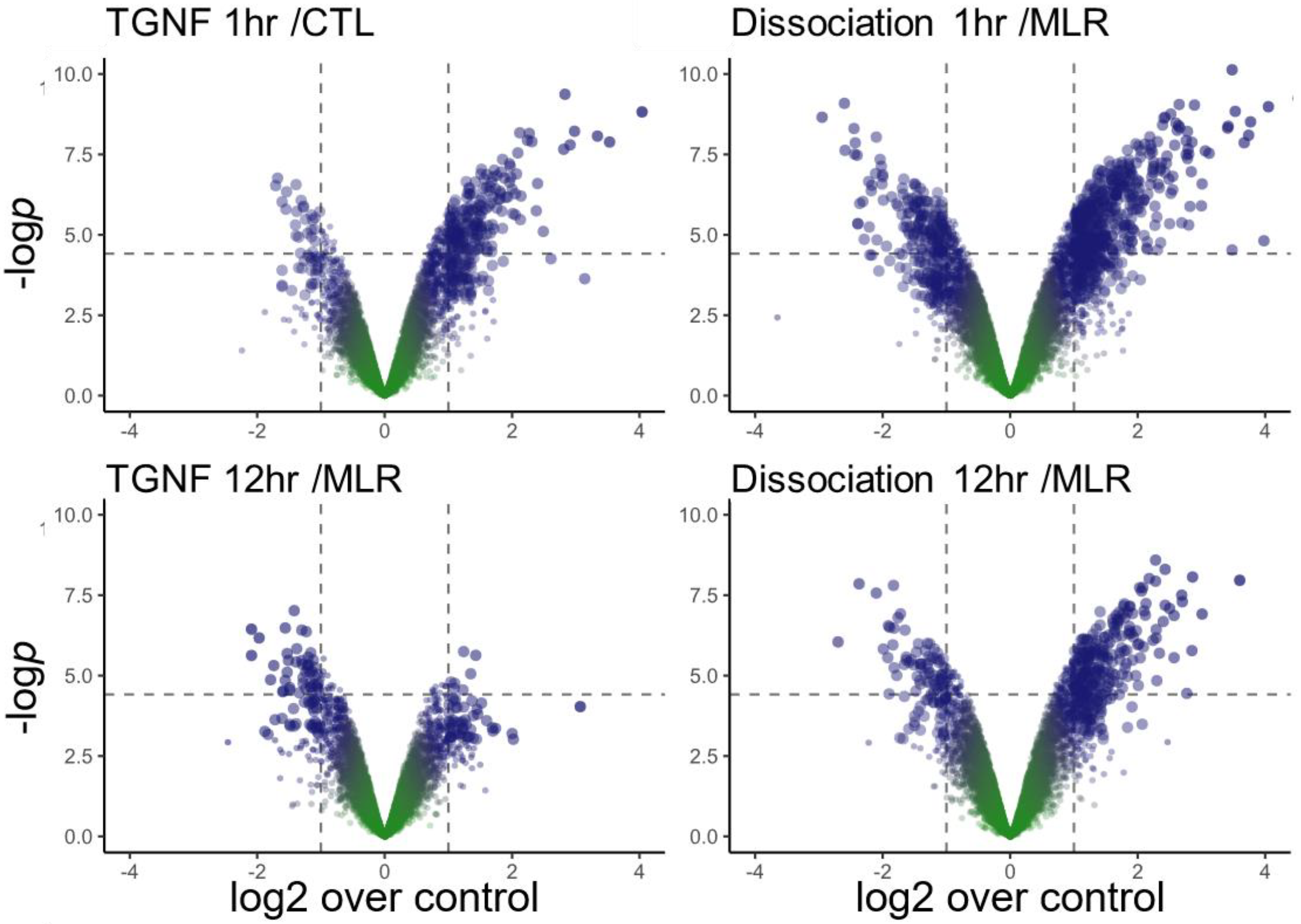
Enzymatic dissociation perturbed many more class I phosphosites, and to a greater degree overall. Volcano plots depicting phosphosite changes over control for each treatment, with dashed lines representing thresholds for significantly changed phosphosites by this method.

**Figure 6.**
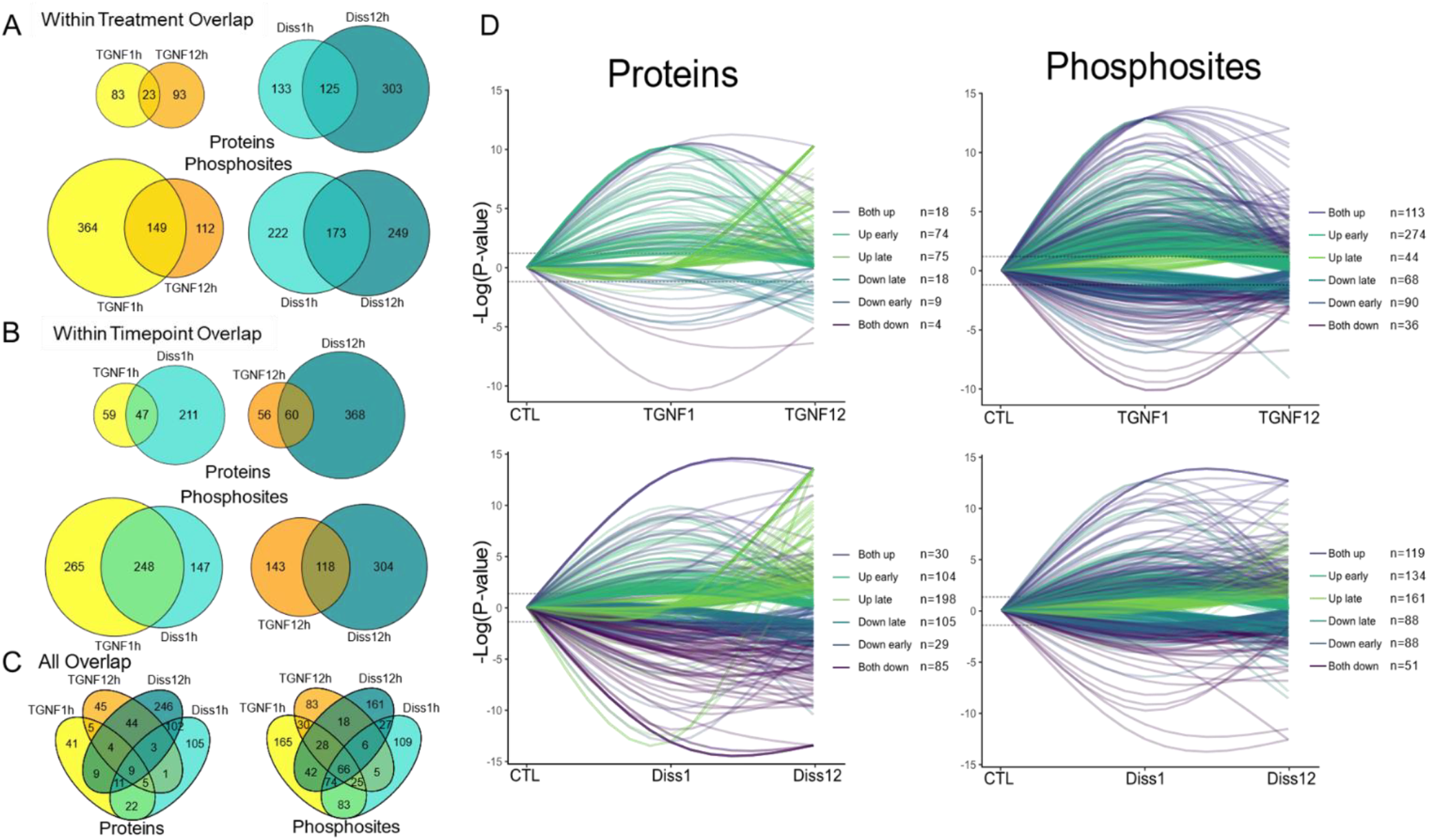
Enzymatic dissociation causes slightly more salient changes than TGNF treatment, but a high proportion of early phosphosite changes are common to both treatments. A-C) A number of protein and phosphosite changes overlap between treatments and timepoints, but the phosphosite changes after 1 hr of both treatments overlapped to the greatest degree. D) Tracking the levels over time for all proteins and phosphosites significantly changed by any condition shows a high proportion of phosphosite changes caused by TGNF at 1 hr that subside by 12 hrs, with other timepoints and treatments showing similar mixes of early, late, and salient changes.

### Signaling pathway enrichment analysis

We evaluated the signaling pathways and biological functions enriched in significantly altered proteins and phosphosites via query against dbEMT, an online database of protein participants in EMT processes (37), and by IPA. Our data included 389 proteins and 260 phosphosites from dbEMT, and 79 were significantly altered. For IPA, the proteins and phosphosites meeting the dual criteria for significance from individual treatments and timepoints were uploaded to the IPA pipeline and the enriched canonical pathways, biological functions, and toxic functions were evaluated.

At 1 hr, TGNF induced phosphorylation changes in *Integrin*, *EGF*, *PDGF*, *Actin cytoskeleton*, *ERK/MAPK*, *HGF*, and *Ephrin B* signaling, among many others, and protein abundance changes in *Cell Cycle Control of Chromosomal Replication*, *Nucleotide excision repair*, *HIPPO*, *CDK5*, and *TNFR1* signaling. IPA was able to detect statistically significant activation (Z-score > 2) in phosphorylation states of *Integrin*, *Actin Cytoskeleton*, *CXCR4*, *Synaptogenesis*, and *Ephrin receptor* signaling pathways. Under the tox functions category *liver hyperplasia* functions – *liver tumor*, *liver carcinoma*, and *liver cancer*-were highly enriched in both the phosphorylation and protein abundance changes with phosphorylation *p*-values of 4.46×10^−20^, 1.88×10^−19^, and 2.22×10^−18^, respectively, and 218 phosphorylated molecules identified as contributing to these functions. The next most highly enriched function was *enlargement of heart* with a 3.46×10^−5^ *p*-value, a gap of 10^13^ from the liver related functions. Meanwhile, *proliferation of hepatocytes* and *proliferation of liver cells* functions exhibited significant activation and several cell death liver-related functions exhibited inhibition. The pathways with the highest combined enrichment from phosphorylation and protein abundance changes were *IL-17A signaling in gastric cells, signaling by Rho Family GTPases, breast cancer regulation by Stathmin1, and EGF signaling.* The most highly enriched canonical pathways in either phosphorylation or protein changes were not among the top enriched pathways in the other, as opposed to the tox functions in which the liver cell proliferation functions were strongly enriched in both.

12 hrs after TGNF treatment, Rho GTPase, epithelial adherens junction, RhoA, Integrin, Actin cytoskeleton, and regulation of Actin-based motility by Rho canonical pathways were significantly enriched in phosphorylation changes along with Wnt/β-catenin, acute phase response, CD40, CD27 in lymphocytes, sertoli cell junction, and hepatic fibrosis/hepatic stellate cell activation pathways in protein abundance changes. Seven pathways were significantly enriched in both the phosphorylation and protein abundance changes caused by TGNF at 12 hrs: Signaling by Rho Family GTPases, Epithelial Adherens Junction signaling, RhoA, ERK/MAPK, Molecular Mechanisms of Cancer, Sertoli Cell-Sertoli Cell Junction signaling, and NGF signaling. Tox functions enriched in 12 hrs changes again included liver cancer and proliferation of hepatocytes, with significant enrichment in both phosphorylation and protein abundance changes.

Dissociation induced 1 hr phosphorylation changes to pathways including Chemokine, Cholecystokynin/Gastrin, AMPK, Aryl Hydrocarbon Receptor, and EGF and protein abundance changes to Hepatic Fibrosis/Hepatic Stellate Cell Activation, Inhibition of Matrix Metalloproteases, FXR/RXR Activation, and LXR/RXR Activation. Thrombin, Synaptic LTP, IL-7, Ephrin B Signaling, and VEGF signaling were significantly activated based on phosphorylation changes and Wnt/β-catenin signaling based on protein abundance changes. Eight canonical pathways were significantly enriched in both phosphorylation and protein changes: Granzyme B Signaling, Aryl Hydrocarbon Receptor Signaling, Agrin Interactions at Neuromuscular Junction, IL-17A Signaling in Gastric Cells, RAR Activation, Tight Junction Signaling, IGF-1 Signaling, and PPAR Signaling. Liver hyperplasia tox functions liver tumor, liver cancer, liver carcinoma again exhibited the strongest enrichment in both phosphorylation and protein changes, along with predicted inhibition of cell death of liver cells in phosphorylation changes.

12 hrs after dissociation, Actin Cytoskeleton Signaling, Tight Junction Signaling, RhoA Signaling, Signaling by Rho Family GTPases, and Integrin Signaling were the most strongly enriched in phosphorylation changes, with Wnt/β-catenin, Hepatic Fibrosis/Hepatic Stellate Cell Activation, HGF Signaling, Role of Macrophages, Fibroblasts & Endothelial Cells in Rheumatoid Arthritis, and LXR/RXR Activation most strongly enriched in protein abundance changes. 47 canonical pathways were enriched in both phosphorylation and protein changes with Tight Junction Signaling, Wnt/β-catenin Signaling, Regulation of Cellular Mechanics by Calpain Protease, Agrin Interactions at Neuromuscular Junction, ILK Signaling, and HGF Signaling topping the combined list. Liver hyperplasia tox functions liver tumor, liver cancer, liver carcinoma once again exhibited the strongest enrichment in both phosphorylation and protein changes.

### Comparison of pathways enriched in different treatments and timepoints

We further sought to examine the overlap between the pathways enriched in proteins and phosphosites altered by the various treatments and timepoints. Correlation analysis of enriched canonical pathways in protein and phosphosite changes of each condition demonstrated overall higher correlation among phosphosite changes, and highest correlation between phosphosite changes observed at the 1 hr timepoints of TGNF and Dissociation (Fig 7).

**Figure 7.**
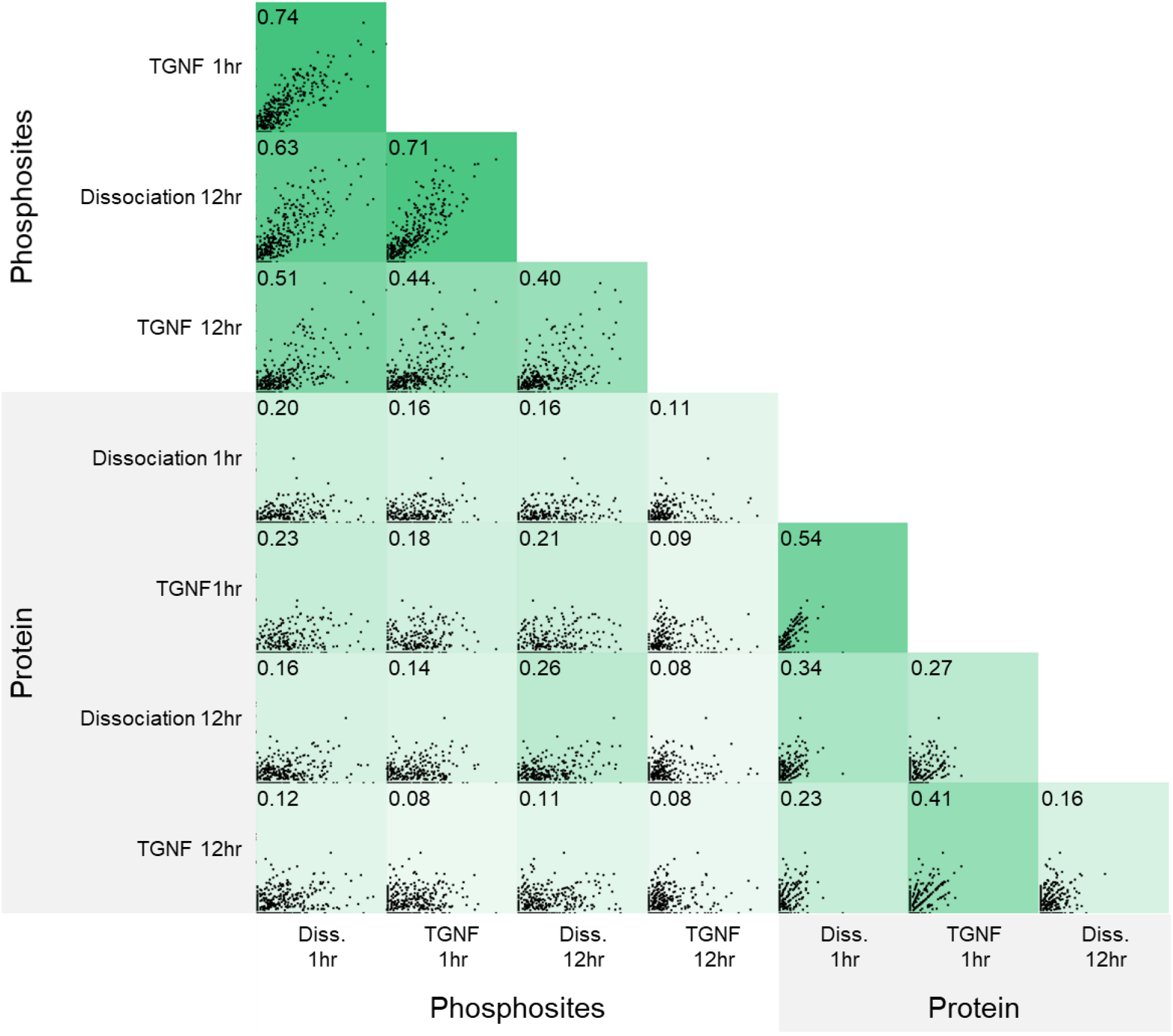
Correlation analysis of the enriched Ingenuity Pathway Analysis (IPA) canonical pathways reveals the 1 hr phosphosite changes for both treatments correlated more strongly than the enriched pathways in any other comparison.

Examining the pathways enriched in combined protein and phosphosite changes showed signaling by Rho Family GTPases, Actin Cytoskeleton Signaling, and Tight Junction signaling exhibited the strongest average canonical pathway enrichment across the 4 conditions, and liver tumor/cancer/carcinoma exhibited the highest average for tox functions (Fig 8). At both TGNF timepoints Signaling by Rho Family GTPases, Integrin Signaling, and EGF signaling were among the most highly enriched, while those changed at both timepoints by dissociation included Hepatic Fibrosis/Hepatic Stellate Cell Activation, LXR/RXR Activation, and Wnt/β-catenin Signaling. Tox functions changed at both timepoints were dominated by liver hyperplasia functions for both treatments, driven most strongly by phosphorylation changes. The signaling pathways enriched in the proteins and phosphosites changed by both treatments at 1 hr included 14-3-3-mediated Signaling, Chemokine Signaling, Apelin Endothelial Signaling Pathway, EGF Signaling, and Signaling by Rho Family GTPases. Liver tox functions including liver cancer and proliferation of hepatocytes were the most strongly enriched in this dataset as well. Overlapping changes at 12 hrs showed enrichment of Integrin Signaling, Tight Junction Signaling, Signaling by Rho Family GTPases, Wnt/β-catenin Signaling, CD40 Signaling, and Cell Cycle: G2/M DNA Damage Checkpoint Regulation. Liver Hyperplasia, Proliferation of liver cells, and Hepatocellular carcinoma were among the most enriched tox functions.

**Figure 8.**
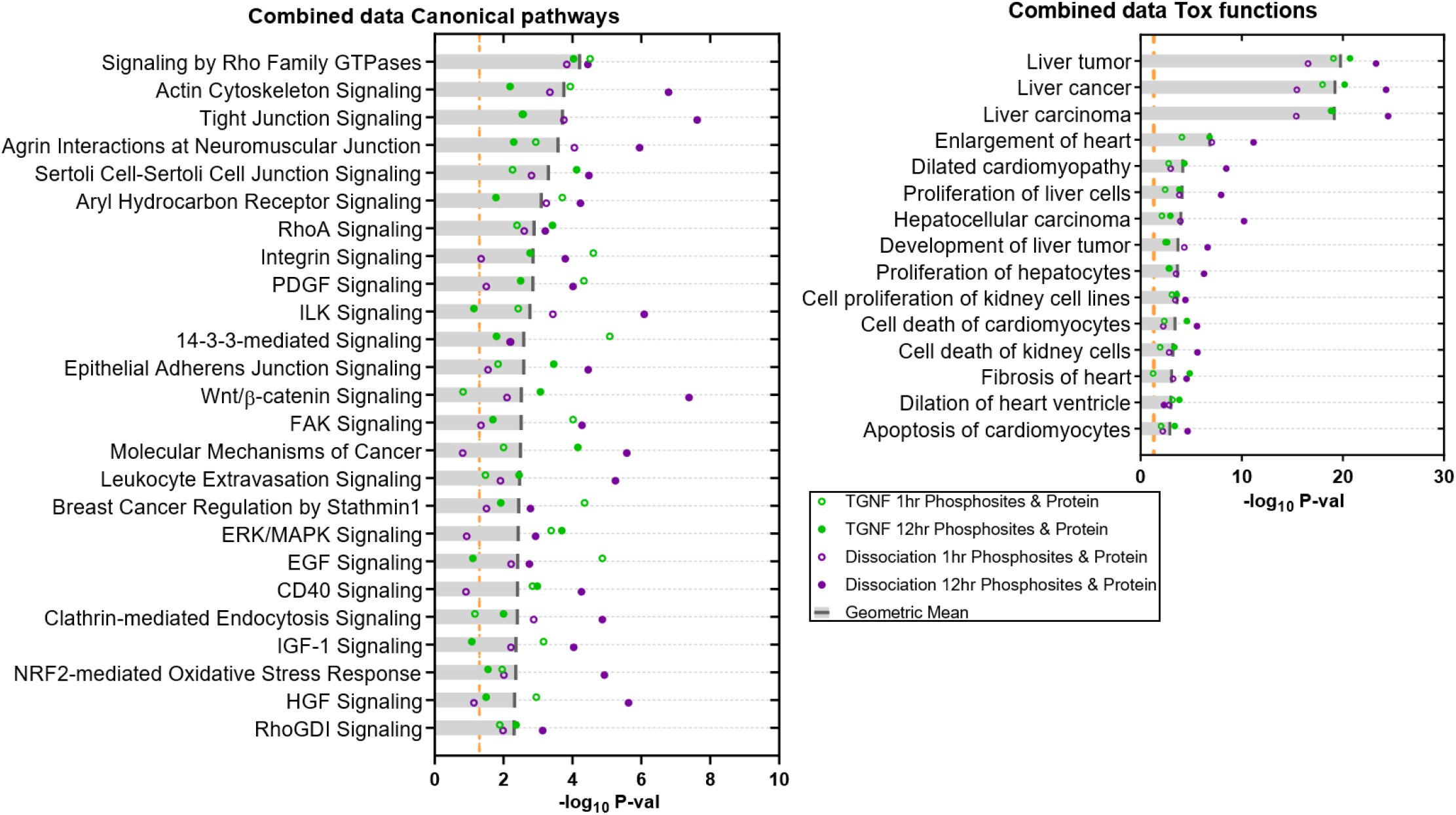
IPA evaluation of the enriched canonical pathways and tox functions in combined protein and phosphosite changes for each treatment reveals a high degree of agreement. Liver tumor/cancer/carcinoma tox functions exhibited extremely high enrichment in the changes caused by all four treatment conditions.

Because phosphorylation changes can occur almost instantaneously while protein changes require more time to develop, we also compared the pathways and functions enriched in both 1 hr phosphorylation changes and 12 hr protein changes for both treatments (Fig 9). 17 pathways were found to be significantly enriched in all four datasets included in this comparison, with the most highly enriched being HGF signaling followed by CD40 Signaling, Sertoli Cell-Sertoli Cell Junction Signaling. Others included Molecular Mechanisms of Cancer, TGF-β Signaling, and Signaling by Rho Family GTPases. HGF (Hepatocyte Growth Factor) signaling being the most highly enriched pathway in this comparison is yet another indication that liver-related signaling is playing an important role under these conditions. Many members of the Liver tumor/cancer/carcinoma tox functions exhibit strong perturbations in protein, phosphosite, or both, and there are many protein-protein interactions between them (Fig 10).

**Figure 9.**
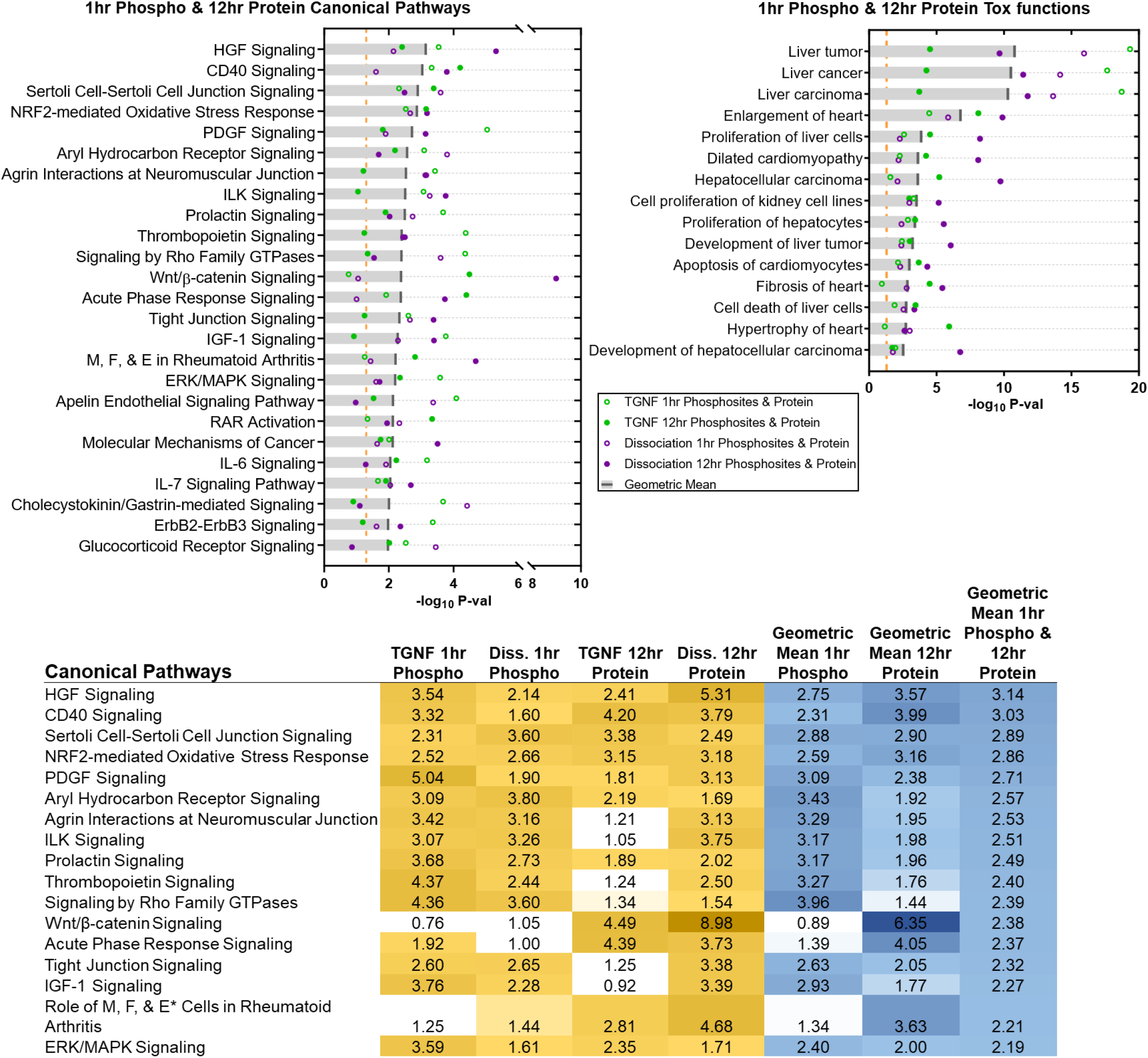
Narrowing the dataset to early (1 hr) phosphoproteome changes and late proteome (12 hrs) changes for each condition reveals the HGF signaling pathway with the strongest combined enrichment among canonical pathways. Liver tumor/cancer/carcinoma tox functions exhibited extremely high enrichment in all the changes caused by all four treatment conditions.

**Figure 10.**
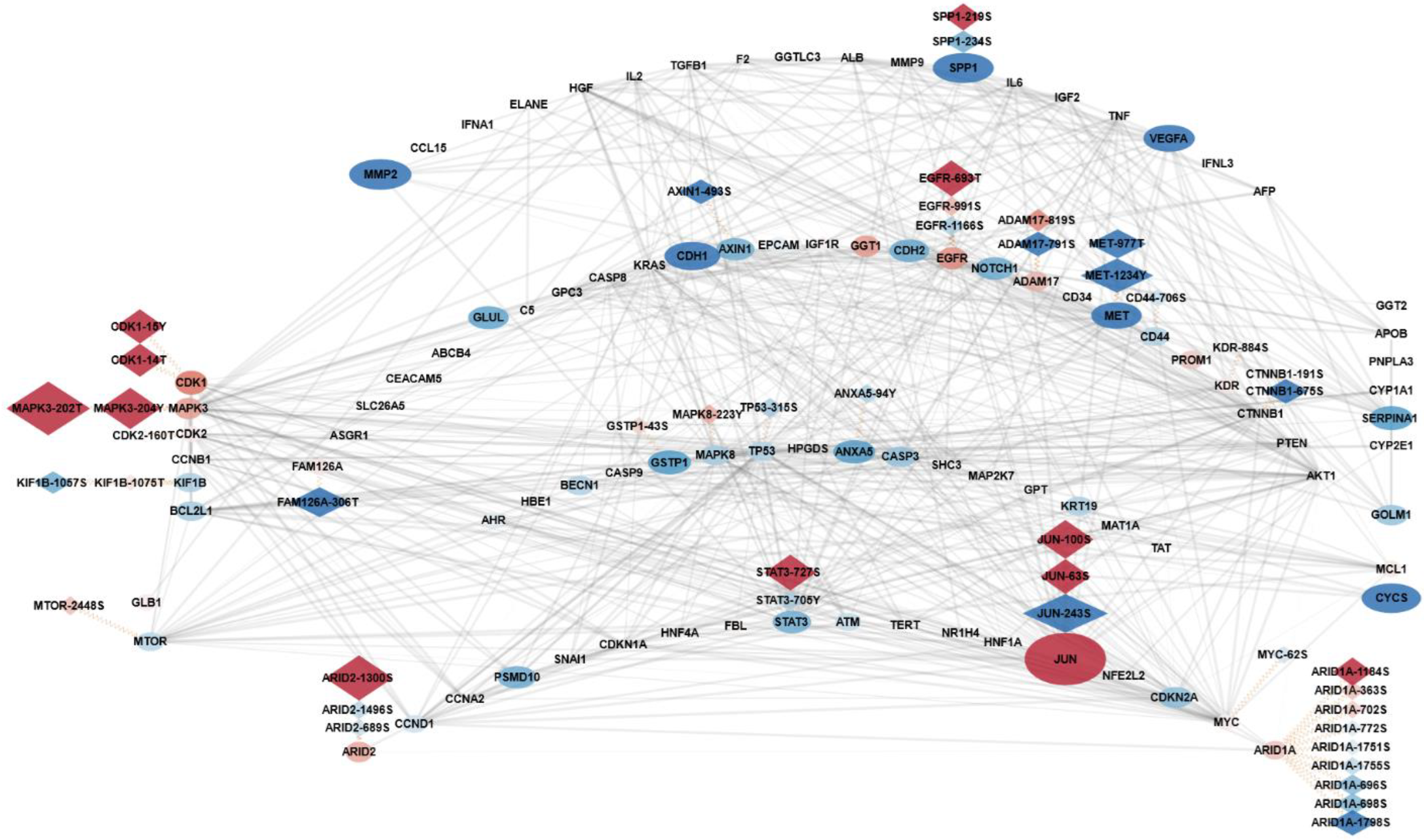
Ingenuity Pathway Analysis revealed numerous perturbations within the “Liver tumor/cancer/carcinoma” tox functions. Both TGNF treatment and dissociation induced strong alterations to many proteins and phosphosites within this biological function, particularly upon the MET cell surface receptor, MAPK3 signaling intermediate, and Jun transcription factor.

### Phosphosite occupancy analysis

To gain a more absolute measure of phosphorylation changes induced by TGNF and dissociation at 1 hr, we calculated phosphosite occupancies. First published by Olsen et al. in 2010 (38), phosphosite occupancy – or stoichiometry – analysis examines the change in abundance of the nonphosphorylated and phosphorylated forms of a peptide caused by treatment and calculates the proportion of that phosphosite that is phosphorylated under both conditions. Several criteria must be met for this analysis, including numerical parameters to allow proper calculations and, importantly, that the peptide be observed in both phosphorylated and nonphosphorylated states. 1063 phosphosites in our data met these criteria (Fig 11A). Both TGNF and dissociation caused an overall increase in occupancy at the sites measured, and while the values of the two treatments correlate well, there was a statistically significantly larger increase caused by dissociation (Fig 11B-E). Intriguingly, the top 5 most strongly shifted phosphosites for both treatments were the same: LMNA-S423, HNRNPK-S116, DMXL1-S924, and EFR3A-S220, all with log_2_FC values over 3, with each exhibiting slightly higher values in response to dissociation. All of these exhibited vanishingly low occupancy in the control state: LMNA-S423 was phosphorylated only 0.06% in the dissociation comparison. It should be noted that occupancy calculations for each treatment produce different occupancy values for the control state through the calculations used, but the difference in the control values for the phosphosites most drastically changed by the two treatments were quite small. Examining raw occupancy changes, HNRNPK-S116 stands out in both treatments with occupancy shifting from ~2% to 53.1% and 80.0% in TGNF and dissociation, respectively. The largest raw decrease was seen in TMSB4X-T34, with a loss of 71.9% occupancy from 93.4% to 21.6% in dissociation, and it saw little change with TGNF treatment. The pathways enriched in the phosphosites with log_2_FC of greater than 1 or less than −1 included Germ Cell-Sertoli Cell Junction Signaling, PAK signaling, Granzyme B Signaling, Integrin Signaling, EGF Signaling and others. Liver hyperplasia/hyperproliferation was the most strongly enriched tox function category for dissociation, but Cardiac dilation, cardiac enlargement showed the strongest enrichment for TGNF.

**Figure 11.**
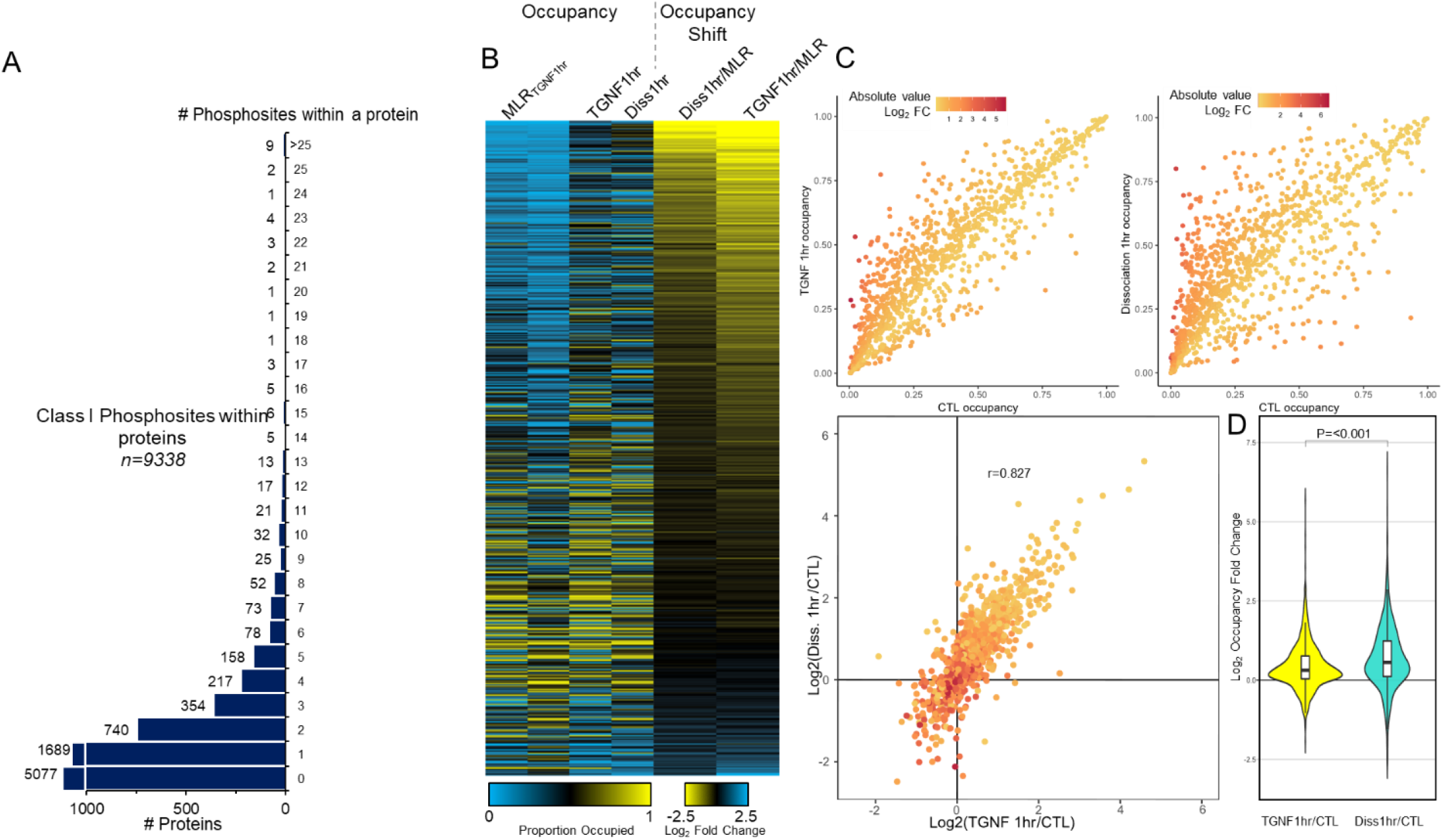
Phosphosite occupancy analysis reveals over 1000 phosphosites with high overall correlation between TGNF and dissociation after 1 hr. A) 1061 phosphosites meet the criteria for phosphorylation occupancy analysis. The vast majority are located within proteins possessing only one phosphosite with occupancy measurements, but 20 phosphosites within the protein SRRM2 were measured. B) Occupancy calculations for the 1 hr timepoints of both treatments reveal similar patterns, particularly in the change from control. C) Plotting the occupancy values for both treatments versus control and versus each other reveal large departures from control and high correlation – but statistically significant difference – between the two (r = 0.827). D) Distribution analysis displays larger gross perturbations caused by dissociation.

### Kinase-substrate relationships

In search of the kinase-substrate relationships (KSR) that have been observed previously between known kinases and the phosphosites in our dataset, we assayed the PhosphositePlus human database updated May 2, 2019. This revealed only 129 phosphosites with known KSRs, paired with 74 kinases. Nine phosphosites in this group (CALD1-789S, FLII-436S, JUN-243S, MAPT-673S, ROCK2-1374S, RTN4-107S, SETMAR-508S, STMN1-25S, and WASF2-308S) surpassed the combined log_2_FC and adjusted *p*-value significance thresholds in at least one treatment, and these matched to twelve kinases. To expand our view we also queried the NetworKIN online database, Version 3.1, downloaded June 6^th^, 2019 (36) to find the kinases that are *predicted* to phosphorylate the sites in our dataset (Fig 12). The algorithm used by NetworKIN calculates kinase-phosphosite phosphorylation probability scores by cross referencing phosphosite peptide sequence versus kinase target motifs, and the connectivity of the two proteins in protein interaction databases. 1,971 phosphosites in our data were found to have at least one kinase relationship of NetworKIN Score > 2.0, with a total of 190 kinases matching with these phosphosites. 237 of these phosphosites surpassed the combined log_2_FC and adjusted *p*-value thresholds in at least one treatment, with 139 significant phosphosites matched to 99 kinases for TGNF at 1 hr, 82 sites and 77 kinases for TGNF at 12 hrs, 99 sites and 101 kinases for dissociation at 1 hr, and 105 sites and 92 kinases for dissociation at 12 hrs.

**Figure 12.**
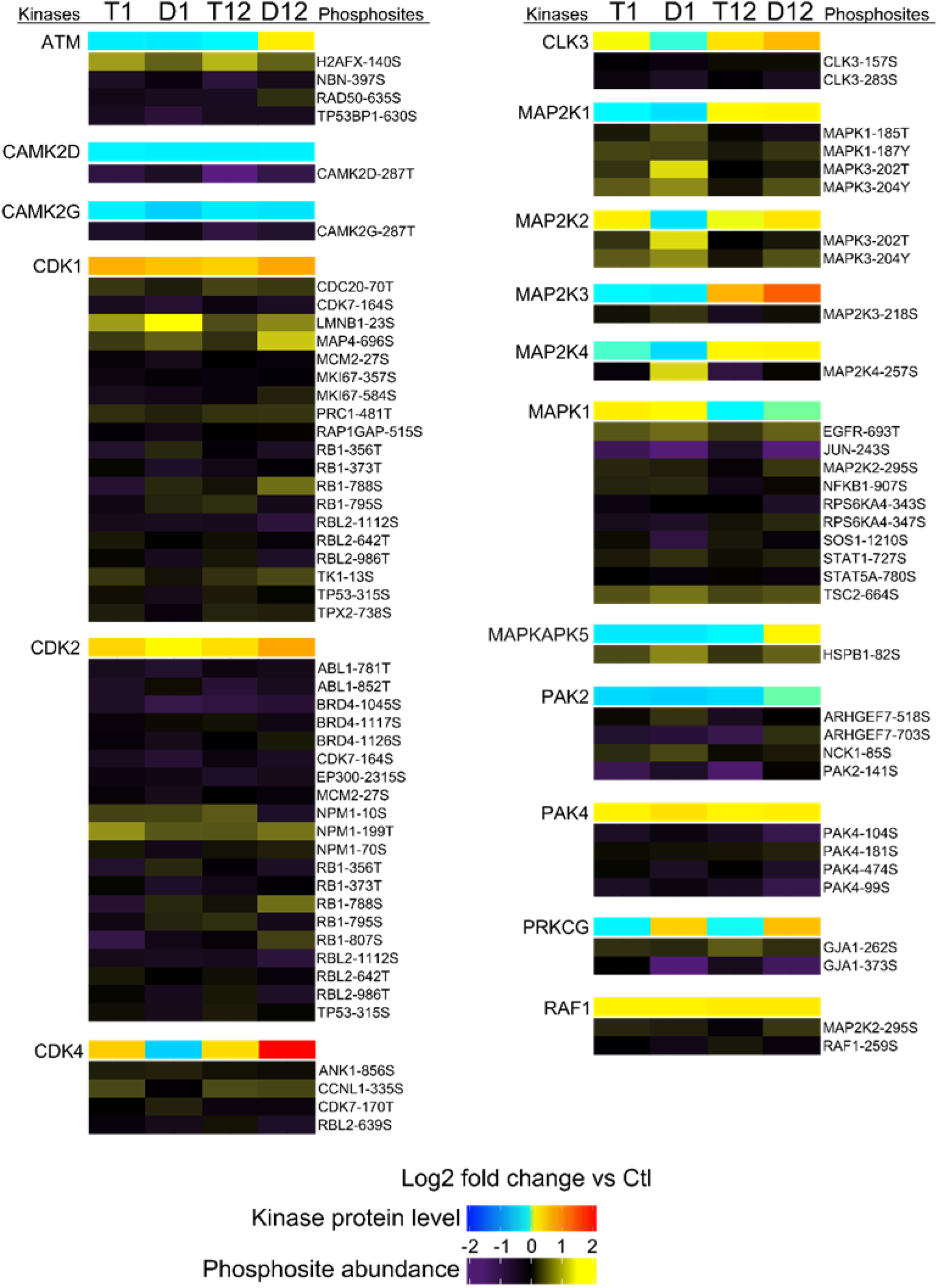
NetworKIN kinase-substrate pair analysis connects many kinases to phosphosites detected in the dataset.

### K-means clustering

As an alternative means of determining meaningful perturbations, we also performed K-means cluster analysis on phosphosite data, choosing a cluster number of 15 as a compromise between avoiding over-clustering and capturing as many meaningful patterns across the groups as possible. Five clusters consisting of 1289 phosphosites that exhibited the largest changes from control were examined for pathway enrichment (Fig 13). The most enriched pathways in these clusters were *FAK Signaling*, *Sertoli Cell-Sertoli Cell Junction Signaling*, *Breast Cancer Regulation by Stathmin1*, *Glucocorticoid Receptor Signaling*, *Tight Junction Signaling*, and *Epithelial Adherens Junction Signaling.* Again, the *Liver Hyperplasia/Hyperproliferation* tox functions exhibited strongly significant enrichment, with *liver carcinoma* topping the list with a *p*-value of 4.07×10^−33^ and 389 molecules involved in this process represented in the five clusters examined.

**Figure 13.**
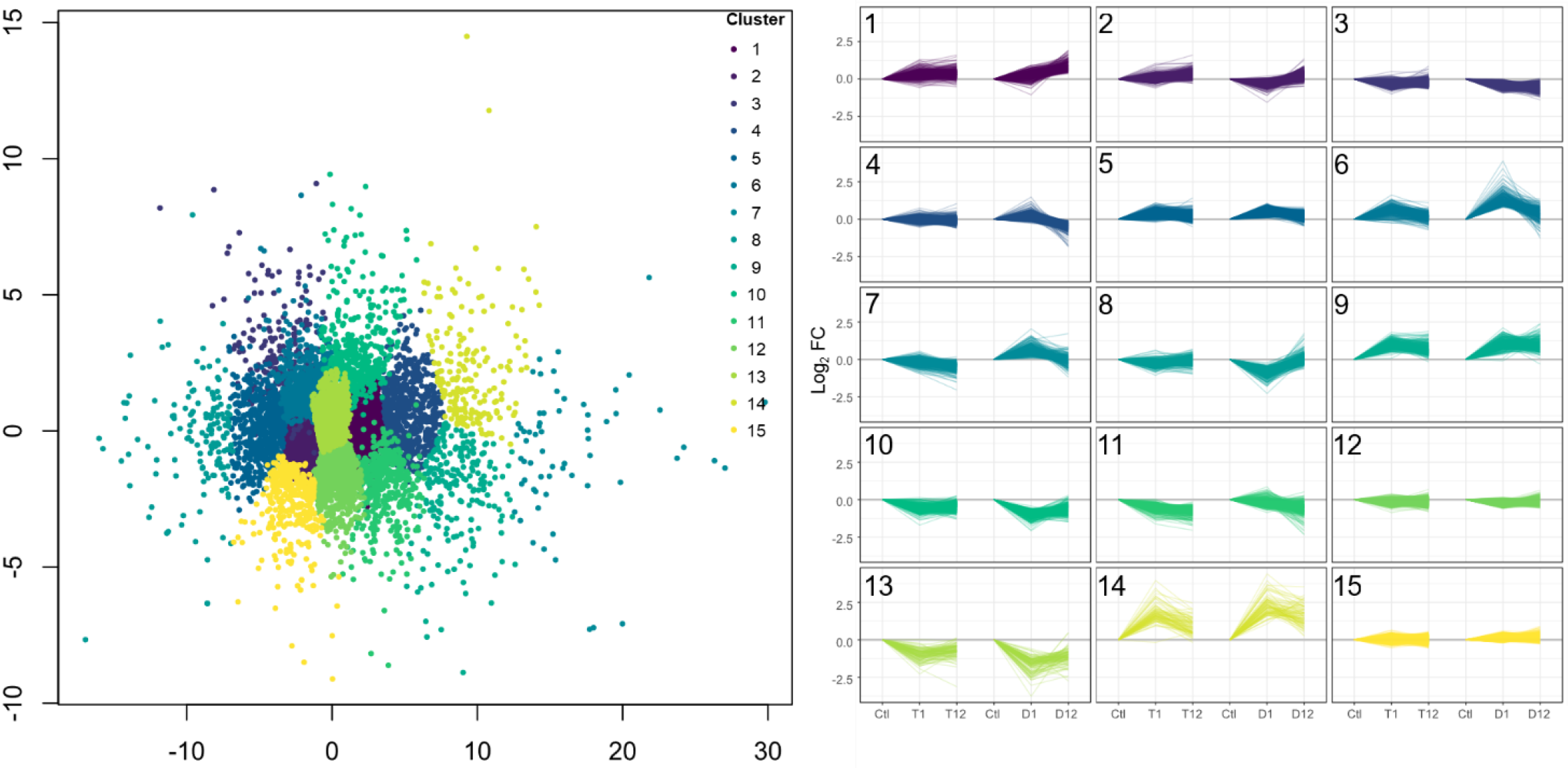
K-means cluster analysis of the phosphoproteome dataset separates several clusters with similar shifts away from control values.

**Figure 14.**
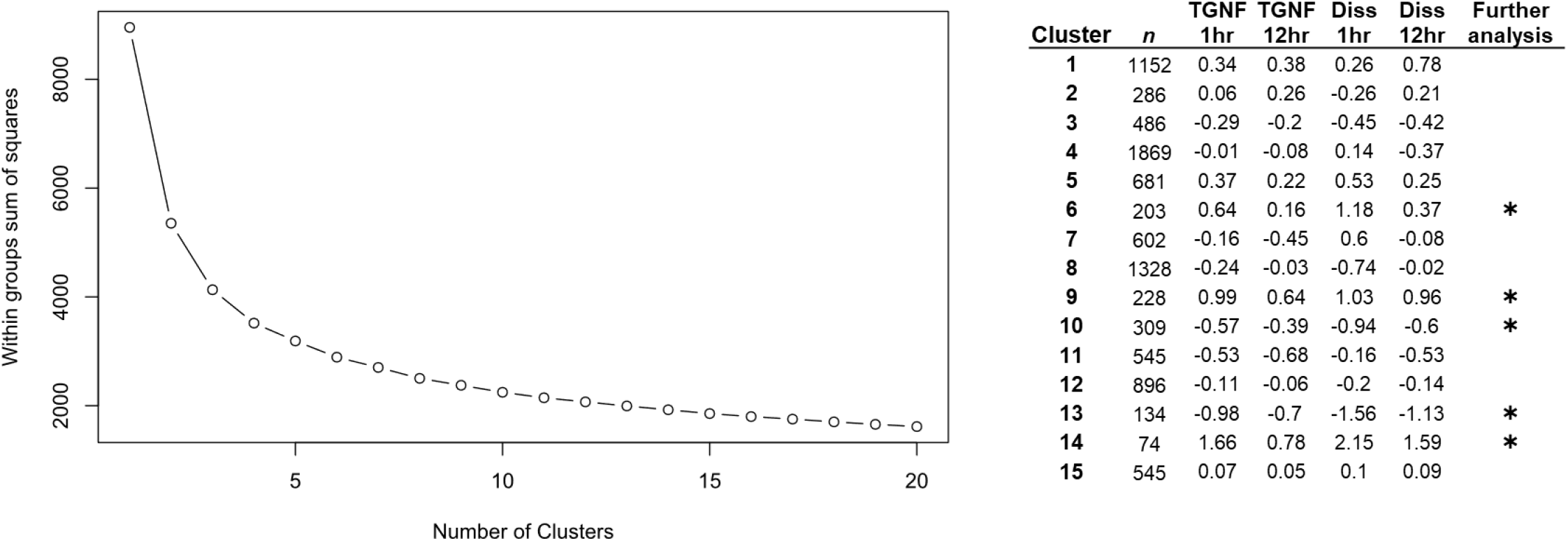
K-means clustering parameters.

## DISCUSSION

A variety of studies have examined RPE protein expression using a variety of methods and cell systems, including immortalized cell lines such as ARPE-19 line (39,40), human donor eyes (41), and more recently RPE derived from human stem cells (42,43). We here add to this prior work by providing extensive proteomic and phosphoproteomic analysis of human stem cell derived RPE, identifying over 8000 total proteins and over 8000 phosphosites quantified across five experimental conditions. The depth we obtained can be attributed to the robust CPTAC proteome/phosphoproteome workflow we employed and to the amenability of the hPSC differentiation protocol to such large-scale analysis. While deep phosphoproteome workflows such as that of CPTAC are often oriented toward abundant sample types such as cell lines and tissues, our RPE differentiation protocol has proved useful for this purpose.

Our data reveal the early stages of EMT protein and kinase signaling in hPSC-RPE cells induced by the well-established combinatorial TGFβ and TNFα treatment (44) and illustrates the contrasts and similarities between TGNF and enzymatic dissociation, a less commonly utilized means of perturbing the epithelial/mesenchymal landscape. The fact that dissociation induces many more protein abundance changes, and more salient phosphorylation changes suggests that enzymatic dissociation is a more widespread perturbation to RPE cellular state. Meanwhile TGNF induces many early phosphorylation changes which then subside likely owing to the fact that it targets several potent kinase cascades, but has longer-term effects that are slower to come online or are more nuanced. The strong overlap in phosphosite changes we observed by both treatments at 1 hr suggests that the two share a large core of signaling alterations. Disruption of cell-cell junctions and loss of cell polarity are key phenomena in EMT, and enzymatic dissociation quickly accomplishes both by cleaving connections both between neighboring cells and between cells and the culture substrate. It seems likely given our observations that doing so activates and inhibits many of the same signaling events affected by TGNF and other well-known EMT induction methods. Thus, it appears dissociation, at least in our hRPE model, effectively bypasses the earliest canonical steps of EMT induction involving secreted growth factor signaling, directly accomplishes some of the major effects of canonical induction and induces many of the same cellular effects. It is therefore likely that disruption of cell-cell connections and cell polarity during EMT aids and/or accelerates the overall EMT process, acting as a positive feedback. It is even possible that it is necessary for continued EMT and might act as a type of checkpoint.

The large alterations we see in canonical liver pathways connect to a body of work that has shown the RPE shares some physiological characteristics of liver. In particular, RPE nearly matches the levels of fatty acid oxidation seen in liver and colon, which exhibit the highest such levels in the body (45). This is thought to be an adaptation to the large amounts of lipids taken in during phagocytosis of photoreceptor outer segments, which the RPE then oxidizes for its own energy demands and is therefore able to transmit more glucose from the vasculature on to the photoreceptors. While our data do not show changes in these specific pathways (i.e. fatty acid oxidation or ketogenesis), the HGF-Met signaling pathway is strongly altered, and it has been observed to regulate these catabolic pathways in other tissues (46). Thus, after only one hour, EMT signaling and/or mechanical dissociation may begin to disrupt the energy supply for the retina, which can very quickly lead to visual deficits.

The capture of many expected changes (cytoskeleton remodeling, adherens junction signaling, EMT signaling, etc.) alongside numerous less expected changes demonstrates the value of deep phosphoproteome analysis. As with any proteomics study, there are also limitations of the data generated. For example, and perhaps surprisingly, we note that several hallmarks of EMT, such as Snai1 and 2, were not observed in our dataset. This is most likely due to the low expression level of these potent transcription factors but could also have been caused by numerous detection limitations inherent to mass spectrometry proteomics, such as those associated with ionization, MS^2^ sampling, and/or limited unique tryptic peptides of appropriate size and composition. The quantitation of thousands of phosphosites across the proteome is a highly valuable set of biological information as kinase signaling pathways are known to transduce and regulate myriad processes. While the functional effects as well as the kinase/phosphatase partners for the vast majority of phosphosites in the RPE are not known, determining the sites that are modulated in a given process, such as EMT, or in response to a given stimulus, can help piece together the complicated kinase networks that regulate and modulate RPE biology, function, and pathology. We hope the data provided by our studies will serve as a useful resource for the RPE research community and help provide the basis for future functional studies of the candidate EMT proteins and phosphosites identified.

## Acknowledgements

The authors would like to thank members of the Zack and Zhang laboratories and Dr. Christina Nemeth for their valuable feedback.

